# Sigma-1 Receptor Promotes Glycolysis in Neuronal Systems by Suppressing GRIM19

**DOI:** 10.1101/2025.07.28.667250

**Authors:** Simon Couly, Yuko Yasui, Ioannis Grammatikakis, Yuriko Kimura, Josh Hinkle, Juan Gomez, Hsiang-en Wu, Ghosh Paritosh, Ashish Lal, Michael Michaelides, Brandon Harvey, Tsung-Ping Su

## Abstract

Sigma-1 receptor (S1R) is a Ca^2+^ sensitive, ligand-operated receptor chaperone protein present on the endoplasmic reticulum (ER) membrane and more specifically at the mitochondria-associated ER membrane (MAM). Upon activation by ER calcium depletion or ligand binding, S1R can increase calcium efflux from the ER into the mitochondria by chaperoning IP3 receptor type3 (Ip3R3). Mitochondrial metabolism has an intricate relationship with glycolysis. Despite S1R affecting mitochondria, the relevance of S1R to glycolysis and its impact on the overall cellular energy metabolism is not known. This study utilizes wild-type (Wt) and S1R knockout (S1R KO) Neuro2a (N2a) cells and Wt and S1R KO mice for primary culture of cortical neurons studies and longitudinal *in-vivo* imaging. In this manuscript we describe the fundamental functions of S1R on glycolysis, mitochondrial activity and NAD^+^/NADH metabolism, keystone coenzymes essential for glycolysis and for mitochondrial activity. Both N2a cells and cortical neurons lacking S1R had reduced glycolytic activity, and increased mitochondria complex I protein GRIM19 but no change in mitochondrial oxygen consumption. Furthermore, we observed an increased NAD^+^/NADH ratio in S1R KO condition. Positron emission tomography revealed decreased [^18^F]fluorodeoxyglucose brain uptake in S1R KO mice. We observed that knocking down GRIM19 in S1R KO condition rescued the glycolysis deficit. Altogether, these data show for the first time that S1R modulates glycolysis and NAD metabolism in various neuronal systems. This new insight on the S1R function may lead to new therapeutic applications of S1R ligands where compromised glycolysis and cellular NAD+/NADH ratios occur such as aging and neurodegeneration.

## Introduction

Sigma-1 receptor (S1R) is a receptor chaperone protein located at the mitochondria-associated endoplasmic reticulum membrane (MAM) [1, 2]. Since its identification in 1982 [3], S1R has been intensively studied across many disease-specific contexts and animal models [4–8]. Several of these studies have demonstrated the relevance of targeting S1R with exogenous ligands in pathological contexts [9–16]; however, the endogenous activation and inactivation of S1R and the downstream consequences are still unclear. At resting state S1R is known to bind with binding immunoglobulin protein (BiP). Upon exogenous ligand activation or endoplasmic reticulum (ER) calcium depletion, S1R translocates along the ER membrane to bind others proteins [2]. One of the most well-known actions of S1R is its binding to inositol 1,4,5-trisphosphate receptor, type 3 (Ip3R3) leading to the stabilization of Ip3R3 [1]. Ip3R3 is a MAM resident protein that facilitates the efflux of calcium from the ER to the mitochondria. Ip3R3 is associated with voltage-dependent anion channel 1 (VDAC1), a protein located on the outer mitochondrial membrane, through a shared interaction with both glucose-regulated protein 75 (GRP75) and translocase of outer mitochondrial membrane protein 70 (TOM70) [17, 18]. Altogether these proteins promote the transfer of calcium from the ER into the mitochondrial intermembrane space. Calcium ions then enter the mitochondrial matrix through mitochondrial calcium uniporter (MCU), a channel protein located in the inner mitochondrial membrane. The increase of calcium inside the mitochondria is complex and downstream consequences vary depending on the intake level and cellular context [1, 17, 19–22].

Mitochondria activity and glycolysis are intricately related [21, 23, 24]. Glycolysis in general is a set of enzymatic reactions that transform molecules of glucose into pyruvate or lactate depending on the oxygen context. However, glycolysis that provides pyruvate to mitochondria is called aerobic glycolysis that favors mitochondrial activity and ATP production from oxidative phosphorylation (OxPhos) when oxygen is available. Conversely, the “anaerobic” glycolysis that leads to lactate without a need for oxygen is conventionally called glycolysis. The OxPhos pathway occurs along five complexes, the first four participate in the electron transfer chain (ETC) and the fifth uses the protons gradient to produce ATP [25, 26]. Although glycolysis and OxPhos occur separately in cytosol and mitochondria, respectively, they both require nicotinamide adenine dinucleotide (NAD) metabolism [27, 28]. In glycolysis, glyceraldehyde-3-phosphate dehydrogenase (GAPDH) and lactate dehydrogenase (LDH) use a form of NAD as substrate. In OxPhos, the complex I, uses NADH, the reduced form of NAD, as substrate to produce NAD^+^, the oxidized counterpart. During this process protons are transferred from the mitochondrial matrix into the intermembrane space to maintain the proton gradient. Mitochondrial activity and glycolysis are further connected through the malate-aspartate shuttle, an active transport process participating in the regulation of NAD^+^ and NADH in the cytosol and the mitochondria matrix [29–31]. Zhang *et al.* and Hou *et al.* provide evidence that downregulating a protein (GRIM19) from complex I of mitochondrial OxPhos promotes glycolysis, highlighting the interplay between the two forms of energy metabolism [32, 33].

From a functional point of view the intricate relationship between mitochondria and glycolysis is observable with metabolic assays [23, 34]. Childers *et al.*, used metabolic assays to observe the oxygen consumption and the acidification rate induced by glycolysis and noted a drastic increase in glycolytic activity when OxPhos complexes III and V were shut down with rotenone and antimycin A [35]. This phenomena is called compensatory glycolysis [34–36] and is observed in different models where mitochondrial activity is inhibited pharmacologically or genetically [37–41].

The relationship between OxPhos and glycolysis has yet another layer of complexity in the brain studies. The brain is one of the highest energy demanding organs with important temporal and spatial fluctuations dependent on the intensity and the nature of the cerebral activity [42, 43]. Another important element in brain related studies is the heterogeneity of the cells that compose the brain and their varying levels of productions and consumptions of energy substrate [44–46].

The impact of S1R on glycolysis, which is known to subsequently alter mitochondria function, has not been observed until recently. In 2020 Motawe et. al [47] used an endothelial cell model to demonstrate that pharmacological activation of S1R enhances glycolysis and decreases mitochondrial oxygen consumption that eventually leads to an overall status quo of the energy metabolism. Oflaz *et al* [48] used two different cancer cell lines to demonstrated that pharmacological activation of S1R increases mitochondrial bioenergetics while decreasing reliance on glycolysis. They also noted that an S1R antagonist did not affect mitochondrial activity but increased glycolysis. These divergent results from Motawe *et al.* and Oflaz *et al.* showcase the complexity of S1R’s role in energy metabolism regulation and could be reconciled given the differences between models and the type of agonist used to activate S1R. However, how would the S1R fashions glycolysis or mitochondrial oxidative phosphorylation in neural system remain unknown.

Here, we investigate the impact of S1R genetic deletion in different neuronal models on glycolysis, mitochondrial activity, and NAD+/NADH metabolism. We found that the S1R elegantly and seemingly very simplistically enhances the neural glycolysis by regulating GRIM19 in the mitochondria remarkably without compromising the mitochondrial oxygen consumption. Results are presented in this report.

## Material and methods

### Cell culture

Neuro2a (N2a) cell line was purchased from American Type Cell Collection (CCL-131). Cells were maintained and grown in Dulbecco’s Modified Eagle’s Medium (DMEM; GIBCO, 11,965–092) containing 10% fetal bovine serum (FBS; R&D Systems #S11150) and 1% penicillin–streptomycin (GIBCO, 15,140–122). Cells were passaged every 2–3 days and maintained at 37 °C in a 5% CO_2_ incubator.

### Generation of S1R KO N2a cell line

Two of the pSpCas9 BB-2A-Puro (PX459) plasmids containing CRISPR guide RNA (gRNA) sequences targeting the mouse sigma-1 receptor (S1R) were obtained from GenScript (SC1948-459). The gRNA sequences, 5⍰-GGCCCCGGG CATAGGCCCGA-3⍰ and 5⍰-CGCTAGAATGCCGTGGGC CG-3⍰ were used. The two plasmids were mixed in 50:50 ratio and transfected into N2a cells. Starting at 48 h after transfection, cells were treated with 2 μg/mL puromycin for 7 days. Cells were trypsinized and resuspended to a density of 8–10 cells/mL. 100μL each of the cell suspension was transferred to a well of a 96-well plate. Expanded cells were collected and cell lysates were analyzed for S1R protein expression by Western blot using the Santa Cruz Biotechnology B5 anti-S1R receptor antibody (sc-137075).

### Transfection

Cell monolayers of 70% density were used for transfection with plasmids using PolyJet reagent (SignaGen Laboratories, SL100688). The PolyJet reagent and plasmids were combined at a ratio of 2:1 in 0.2 mL serum-free DMEM and incubated for 12 minutes at room temperature. Subsequently, mixed DNA-PolyJet complexes were added into each well. Media was refreshed after overnight incubation. Cells were then incubated at 37 °C in a 5% CO_2_ incubator (Thermo Fisher Scientific) for 48 hours after transfection. For stable transfection, the protocol was carried out according to the stable transfection protocol described by ThermoFisher Scientific. Plasmids used are presented in Appendix 1.

### Short hairpin RNA

Short hairpin (Sh) RNA for GRIM19 and scramble control were obtained from by Sigma-Aldrich MISSION®. We ordered custom cloning of two TRC clones and one nontargeting control into the (SHV14) pLKO.1_U6-shRNA_hPGK-NEO_CMV-tGFP vector. Sigma-Aldrich produced plasmid DNA TRCN0000287887; TRCN0000126394; NTC target sequence: 5’-GGCGGGCCATTTGTTTCAATAT-3’; Formulation/Fill Volumes: 1ug of plasmid DNA, 50ul at 20ng/ul.

### Primary culture of cortical neurons

The day before dissection, dishes or round coverslips were coated overnight with phosphate buffered saline (PBS) containing Poly-D-lysine (PDL) at 0.1mg/mL (Sigma-Aldrich, P6407) at 4°C. On the day of dissection, the coverslips were rinsed three times with PBS and then air-dried until use. Pre-culture media and culture media solutions were prepared prior to dissection. Pre-culture media contained Neurobasal Plus (Gibco, A3582901); 10% FBS (R&D Systems, S11150); 1% Penicillin-Streptomycin (Gibco, 15140122); and 1% GlutaMAX (Gibco, 35050061). Culture media contained Neurobasal plus; 2% B-27 Plus (Gibco, A3582801), 1% Penicillin-Streptomycin; 1% GlutaMAX. We also prepared solutions of 2.5% trypsin (Sigma-Aldrich, T4799-25G) resuspended in PBS and 50mg/mL DNase (Sigma-Aldrich, DN25). All solutions were maintained at 37°C except for DNase which was kept on ice.

Pregnant mice were quickly sacrificed, and embryos were extracted. Brains were isolated from embryos and submerged in cold PBS. Brain cortices were isolated under a binocular microscope, cut in small pieces, and placed in tube with 18mL of ice-cold PBS. Under a sterile hood 2mL of trypsin and 200µL of DNase were added to the tube. Tubes were then placed in an incubator at 37°C and mixed frequently for 15min. 20mL of pre-culture media was added to the tubes to stop the trypsin reaction. After pelletizing cells with a 3-minute centrifugation at 600g, the pellet was resuspended gently with 10mL of pre-culture media and then filtered through a 100µm nylon mesh. Centrifugation and resuspension were then repeated twice more. Cells were then counted using a trypan blue solution and hemacytometer and seeded in pre-culture media. After three hours pre-culture media was replaced with culture media. Cells were then kept in a 5% CO_2_ incubator at 37°C for 6 days and media was replaced every other day.

### Immunofluorescence and confocal microscopy

Cells grown on coverslips were washed three times with PBS then fixed for 20 minutes with PBS containing 4% paraformaldehyde (PFA). Cells were then rinsed three times with PBS before mounting a coverslip with 3µL of Dapi-Fluoromount-G™ (Electron Microscopy Sciences, ref17984-24). Pictures were taken with Zeiss Confocal laser scanning microscopes LSM710 or with Olympus CKX53 with CoolLed system. GFP-positive cells and DAPI positive cells were counted and transfection efficiency was calculated as a number of GFP-positive cells/number of DAPI-positive cells.

### Protein extraction

Forty-eight hours after transfection, cells were washed with PBS and harvested by scraping into 1.5 mL tubes. Tubes were then centrifuged at 600g for 3 minutes and the pellets were lysed with 100 μL of modified radioimmunoprecipitation assay (RIPA) buffer (50 mM Tris–HCl, pH 7.4, 150 mM NaCl, 0.05% sodium dodecyl sulfate [SDS], 0.5% Triton X-100, and 0.05% sodium deoxycholate [SigmaAldrich, D6750]) supplemented with ethylenediaminetetraacetic acid (EDTA)-free protease inhibitor cocktail tablets (Complete, Mini, EDTA-free; Roche Diagnostics, 11,836,170,001) and phosphatase inhibitor cocktail tablets (PhosStop, Millipore, PHOSS-RO). Tubes were incubated on ice for 10–15 minutes and then centrifuged for 10 minutes at 18000 g and 4 °C; the supernatant was collected and stored at −80 °C for protein analysis.

### Western Blot

Protein concentrations of cell lysates were determined using BCA protein assay kit (Thermo Fisher Scientific, 23,225). 4X Laemmli sample buffer (Bio-Rad, 161–0747) containing 10% 2-mercaptoethanol was added to 30 μg of protein. Samples were heated for 5 minutes at 90 °C. Protein samples were then separated by using 14% SDS–polyacrylamide gel electrophoresis (SDS-PAGE) containing 2% of 2,2,2-trichloroethanol to visualize the quantity of total protein in each lane and transferred overnight at 4°C onto a polyvinylidene fluoride (PVDF) membrane at 30mA. After incubation with 2% bovine serum albumin (BSA) in tris-buffered saline with 0.1% Tween 20 (TBST; Bio-Rad Laboratories, 170–6531) for 1 hour, membranes were incubated with primary antibodies (see Appendix 1) for 1 hour at room temperature. Membranes were washed three times with TBST for 10 minutes, followed by an incubation with secondary antibodies (see appendix 1) for 1 hour at room temperature. Blots were washed three times for 10 minutes with TBST and developed using the Li-Cor Odyssey CLx. Band intensity was analyzed by Image Studio Lite (LiCor 5.2.5) according to the manufacturer’s manual. Signal intensities were normalized to total protein quantified with 2% of 2,2,2-trichloroethanol.

### RNA extraction, cDNA synthesis and RT-qPCR

Total RNA was extracted using TRIzol (Invitrogen) according to the manufacturer’s protocol. For cDNA synthesis, we used iScript reverse transcription supermix (Bio-Rad) with 500 ng of the extracted RNA. For real-time qPCR, all reactions were carried out on StepOnePlus real-time PCR System (Applied Biosystems) using FastStart SYBR Green Master Mix (Millipore Sigma). Glyceraldehyde-3-phosphate dehydrogenase (Gapdh) mRNA was used to normalize expression, and the relative expression of RNA was calculated using the threshold cycle (2−ΔΔCT) method. RT-qPCR primer sequences for each gene are indicated as follows:

Gapdh: AACTTTGGCATTGTGGAAGG CACATTGGGGGTAGGAACAC
Grim19: CATCGGGGCCTTGATCTTTG GGCCTCCAAGTCCTCAATCA

### Polysome fractionation

N2a Wt and S1R KO cells were grown in 10cm dishes to ∼90% confluency and then treated with 100⍰μg/mL cycloheximide (Sigma-Alrich) for 10 minutes. Cytoplasmic lysates were prepared using Polysome extraction buffer (100⍰mM KCl, 5⍰mM MgCl2, 0.5% NP-40, 100⍰μg/mL cycloheximide, 2⍰mM dichlorodiphenyltrichloroethane [DDT], 40 U/mL RNase Out [Invitrogen], and 1× complete protease inhibitor cocktail). Lysates were then loaded in a 10% to 50% linear sucrose gradient and then fractionated by ultracentrifugation in an SW 40 Ti swinging bucket rotor (Beckman Coulter) at 37,000⍰rpm at 4°C for 2 hour. Twelve fractions were obtained via a fraction collector (BioComp Instruments) and 250 μL from each fraction was used to extract RNA using TRIzol LS (Thermo Fisher). From each fraction, equal volumes of RNA were used for cDNA synthesis and reverse transcription quantitative real-time polymerase chain reaction (RT-qPCR).

### Metabolic flux assays

N2a cells were plated with their specific growth media at a density of 6×10⁴ or 2×10^4^ cells per well in their specific growth media within respectively Seahorse XF24 or XF96 Cell Culture Microplate, respectively. For the primary culture of cortical neurons, plates were precoated with PBS containing poly-D-lysine (PDL) at 0.1 mg/mL overnight at 4°C and seeded at a density of 5×10^5^ or 2×10^5^ in XF24 or XF96, respectively. For both N2a cells and cortical neurons, on the day of the culture PDL was removed, and wells were washed three times with PBS and air dried until used. The plate was gently rocked to ensure even cell distribution before incubating at 37⍰°C overnight to allow cell attachment. N2a transfections, media was removed the following day and replaced with transfection reagent for 24 hour, followed by normal media replacement and subsequent Seahorse assays. For the primary culture of cortical neurons, cells were tested after seven days *in-vitro*. On the day of the experiment for both N2a cells and cortical neurons, the growth media was replaced with Seahorse Phenol Red-free DMEM, and oxygen consumption rate (OCR) and extracellular acidification rate (ECAR) were measured with the Agilent Seahorse XF Cell Mito stress test and Glycolytic Rate assay, respectively. Assays were carried out according to the manufacturer’s guidelines. After completing the XF24 Seahorse assay, cells were collected and processed for BCA assay to normalized seahorse results with the protein quantity contained in each well. For XF96 assays, plates were scanned with Celigo Image Cytometer to normalized seahorse results with number of cells in each well.

### PET-scan

For PET-scan longitudinal analysis we observed the evolution of the uptake of [18F]FDG (fluorodeoxyglucose) in the brain of 8 Wt and 8 S1R-KO male mice at different ages : 2, 3, 6, 9 and 12 months. For each time point, mice were fasted overnight prior to the experiment. The next day, the mice received an intraperitoneal injection of [18F]FDG (Cardinal Health) and placed back into their home cages. After 30 minutes, the mice were anesthetized with 5% isoflurane, placed on a custom-made bed maintained under anesthesia (1-2% isoflurane), and scanned with a nanoScan© small animal PET/CT scanner (Mediso) for 20 minutes. A CT scan was acquired at the end of the PET scan, and the mice were returned to their home cage. In all cases, the PET data were reconstructed using the nanoScan© built-in algorithm, correcting for attenuation and radioactive decay with a voxel size of 0.4 mm. Images were coregistered to an MRI template using PMOD software (PMOD Technologies) and then analyzed using MATLAB R2023 (MathWorks) and SPM12 (University College London). Voxel-based repeated-measures Student’s t tests were performed, and the resulting parametric images were filtered for statistically significant (P < 0.05) clusters larger than 100 contiguous voxels. Additionally, VOI values corresponding to the frontal cortex, dorsal striatum, and basal forebrain were drawn using a PMOD template. The VOI values (kBq/cc) were extracted, Standard Uptake Value (SUV) was calculated by adjusting for body weight and injected radiotracer (µCi).

### NAD^+^ /NADH measurement

NAD^+^ /NADH assays were performed with NAD/NADH-Glo™ Assay (Promega, refG9071) on N2a cells and primary cortical neurons in 96 well plates. N2a cells and cortical neurons were seeded at 2×10^4^ and 2×10^5^ cells per well, respectively. For N2a cells, assay was carried out 48 hours after seeding or 48 hours after transfection. For primary culture of cortical neurons, assay was carried out after 6 days *in-vitro* (DIV). Assay was carried out according to the manufacturer’s guidelines.

### Peredox assay

N2a cells were seeded in µ-Slide I0.4 Luer ibiTreat (ibidi, 80176) at a density of 4.5×10^4^ cells per slide. The next day, cells were transfected with pMOS023: Peredox NADH/NAD+ cytosolic sensor (a gift from Adam Cohen; Addgene plasmid # 163060; http://n2t.net/addgene:163060 ; RRID:Addgene_163060) [55]. The following day the transfection media was replaced with culture media. Forty-eight hours after transfection, the cells were incubated in media containing 121.5mM NaCl, 25mM NaHCO3, 2.5mM KCl, 2mM CaCl2, 1.25mM NaH2PO4 and 1mM MgCl2. Before the experiment, different preparations of this media were made with varying concentrations of Sodium-L-Lactate (Sigma Aldrich, 71718-10g) and Sodium Pyruvate (Sigma Aldrich, P2256-25g), as described in Appendix 2. Cells were then imaged using a Nikon Eclipse Ti2 A1R inverted confocal microscope (20x Plan Apo VC DIC NA 0.75 objective; Nikon Instruments, Japan) with a temperature-and gas-controlled chamber (37°C; Tokai Hit incubation system, WSKM GM8000, Japan). Ibidi chambers with cells were superfused with fresh media via peristaltic pump (MasterFlex Ismatec Digital MiniFlex, MFLX7801840, Germany). In each instance of lactate/pyruvate media change, the pump was stopped and restarted. T-Sapphire “green” signal was acquired by exciting the sample at 405nm and mCherry “red” signal was acquired by exciting the sample at 561nm (the 488nm laser remained off). Background subtraction and analysis of the change in green/red ratio over time [49, 50] were performed with Nikon NIS Elements software (AR v5.2.04, Japan).

### Glucose uptake assay

Glucose uptake assays were performed using Glucose Uptake-Glo™ Assay (Promega; ref J1342) on N2a cells and primary cortical neurons in 96 well plates. Cells were seeded at 2×10^4^ and 2×10^5^ cells per well, respectively. N2a cells were assayed after 3 days of seeding, and primary culture of neurons were assayed at 6 DIV. Assays were carried out according to the manufacturer’s guidelines. After the assays were completed, N2a cells were processed for a BCA assay to normalized Seahorse results to the protein quantity in the well. Results from primary neuronal culture results were normalized to the number of cells.

### Animals

All methods and animal procedures were conducted in accordance with the principles as indicated by the NIH Guide for the Care and Use of Laboratory Animals. These animal protocols were reviewed and approved by the NIDA intramural research program Animal Care and Use Committee, National Institute of Health. Male and female, Wt and S1R KO mice were used in this study. S1R KO were generated as previously described [51]. We obtained the mice of Oprs1 mutant (+ / −) OprsGt(IRESBetageo)33Lex litters on a C57BL/6 J × 129 s/SvEv mixed background from the Mutant Mouse Regional Resource Center of the University of California, Davis. The S1R (+ / −) males were back-crossed for 10 generations to females on C57BL/6 J and then, mice were further genotyped. Mice were maintained in a 12 h day/night cycle facility with free access to food and water.

### Statistics

All statistical analyses were performed with Prism V10.4.1. All graphs present SEM and individual data points (circles) over mean (bar).

## Results

### Knocking out S1R decreases glycolysis but did not change oxygen consumption in N2a Cells

Previous studies observing the impact of inactivation or deletion of S1R used pharmacological (antagonist) or genetic tools (shRNA and siRNA), allowing only observations of the short-term consequences of a partial S1R inactivation. In this study we mutated the genomic S1R to observe the consequences of S1R knockout in mammalian neuronal models. Thus, we knocked out S1R from Neuro-2a cells (N2a) neuroblastoma via CRISPR-Cas9 (Figure 1A). We observed the energy metabolism using Seahorse, which measures local pH (extra-cellular acidification rate or ECAR) that depend on the proton efflux rate (PER) from the cells. Simultaneously to the ECAR measurement, Seahorse also measures the oxygen consumption changes (oxygen consumption rate or OCR) at the vicinity of the cells. These measurements are dependent on glycolytic and mitochondrial activity but not only. To accurately measure the PER due to glycolysis (glycoPER) and the OCR due to mitochondria (mitoOCR) molecular inhibitors of these pathways are used. To measure glycoPER, we applied a sequence of drugs: First a mix of Rotenone and Antimycin A (Rot/AA) then 2-deoxyglucose (2-DG). Rotenone and Antimycin A (Rot/AA) block mitochondrial-mediated extracellular acidification and then 2-deoxyglucose (2-DG) allows to measure others non-glycolysis acidification.

**Fig 1.**
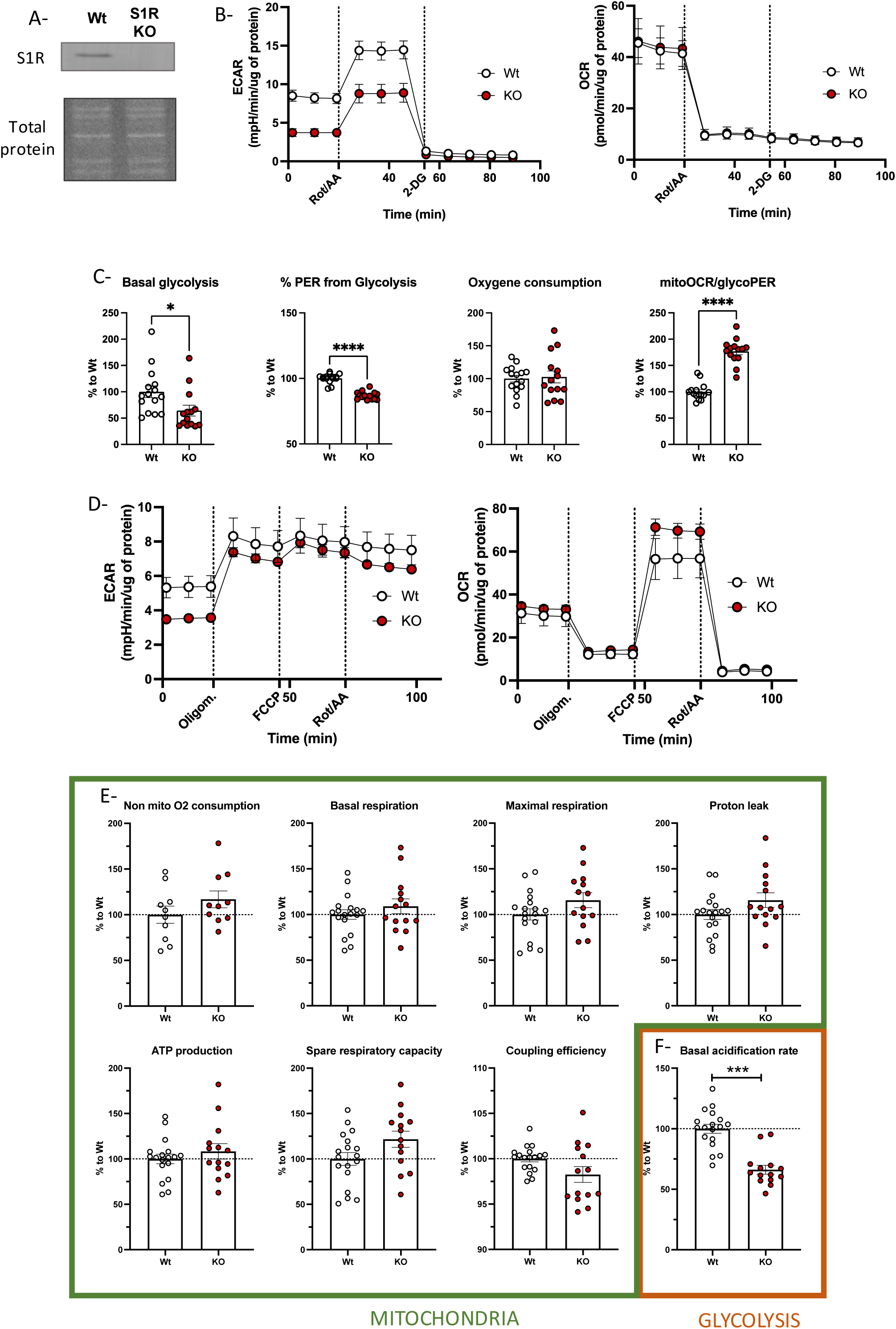
Glycolysis activity and mitochondria Respiration in Wt N2a and S1R KO N2a cells. A-Western blot analysis of S1R expression and total protein staining in N2a cell protein extract. B (left)-N2a cell extra-cellular acidification rate (ECAR) measurement with the application of Rotenone and Antimycin A (Rot/AA) and 2-Deoxyglucose (2-DG); (right)-Simultaneous measurement of N2a cell oxygen consumption rate (OCR). C (from left to right)- Basal glycolysis; % proton efflux rate (PER) from glycolysis; Basal oxygen consumption; MitoOCR/glycoPER. D (left)-N2a cell extra-cellular acidification rate (ECAR) measurement with the application of Oligomycin (Oligom.), Carbonyl cyanide-p-trifluoromethoxyphenylhydrazone (FCCP) and Rotenone and Antimycin A (Rot/AA); (right)-Simultaneous measurement of N2a cells oxygen consumption rate (OCR); E (from left to right)-Non mitochondrial oxygen consumption; Basal respiration; Maximal respiration; Proton leak; ATP production; Spare respiratory capacity; Coupling efficiency; Basal acidification rate. Statistical analysis: Student’s t-test (*p < 0.05,***p < 0.001). B-E Wt N2a in white, S1R KO N2a in red.

We observed a lower ECAR in S1R knockout (KO) cells compared to wildtype (Wt), despite similar OCR (Figure 1B). There was also a significant decrease in basal glycolysis and percentage of PER due to glycolysis (%PER) in S1R KO cells compared to that of Wt; however, there was no difference in basal oxygen consumption between the two cell types (Figure 1C). This led to an increased ratio of mitoOCR/glycoPER in S1R KO cells (Figure 1C). We did not observe any changes in OCR, which is a raw estimation of mitochondria activity, under any conditions. To confirm this result, we then focused on mitochondrial measurement. We used Oligomycin and Rot/AA to calculate the amount of oxygen consumed by mitochondria and used FCCP (carbonyl cyanide p-(trifluoromethoxy)phenylhydrazone) to observe mitochondrial maximum capacity (Figure 1D-F). Consistent with Rot/AA and 2-DG measurements, the ECAR was decreased in S1R KO cells compared to that of Wt, without a corresponding change in OCR (Figure 1D). Furthermore, we did not observe changes is other parameters indicative of changes in mitochondrial metabolism (Figure 1E); only the basal acidification rate was significantly reduced in S1R KO cells compared to that of Wt, which reflects decreased glycolytic activity (Figure 1F). These results correlate with data previously published [52] showing the pharmacological manipulation of S1R does not change the basal mitochondria respiration. However, for the first time we report that a deletion of S1R induces a change in glycolytic activity.

### Knocking out S1R affects glycolytic machinery and complex I protein in N2a cells

We hypothesized that the functional defects in glycolysis could be caused by changes in glycolytic enzyme concentrations. Therefore, we measured the protein concentration of hexokinase (Figure 2A), pyruvate kinase (PKM) (Figure 2B), GAPDH (Figure 2C) and enolase (Figure 2D) in protein extracts from Wt and S1R KO N2a cells. We observed a significant decrease in enolase concentration in the KO condition compared to that of Wt. We did not note statistically significant changes in levels of hexokinase, PKM and GAPDH. In addition, we measured the concentration of proteins belonging to complexes, I, II, III, IV and V of the oxidative phosphorylation (OxPhos) chain; that is, GRIM19, SDHA, UQRC2, MTCO1, ATP5a, respectively. We found no change in protein level in complexes II (Figure 2F), III (Figure2G), IV (Figure 2H) and V (Figure 2I). However, we observed a significant increase in GRIM19 concentration (Figure 2E). This result is correlated with results from Abdullah *et al.* [13] demonstrating that in cardiomyocytes from S1R KO animals have increased complex I protein concentration and no changes in the proteins belonging to other OxPhos complex at first. Another group [52] described no effect on respiration when using S1R ligands but noted an increase of complex I activity with the use of S1R agonist in a Ca2+-dependent manner. However, the same group did not observe any change in the activity of the others OxPhos complexes. It is well known and frequently reviewed [53] that among the OxPhos chain, complex I is not directly involved in the oxygen consumption (complex IV), nor in ATP production (complex V) but participates in the regulation of the NAD^+^ /NADH system as it use NADH as a coenzyme. This could explain why we did not observe any change in the OxPhos activity despite the increase in a complex I protein GRIM19, using an oxygen consumption-based functional measurement.

**Fig 2.**
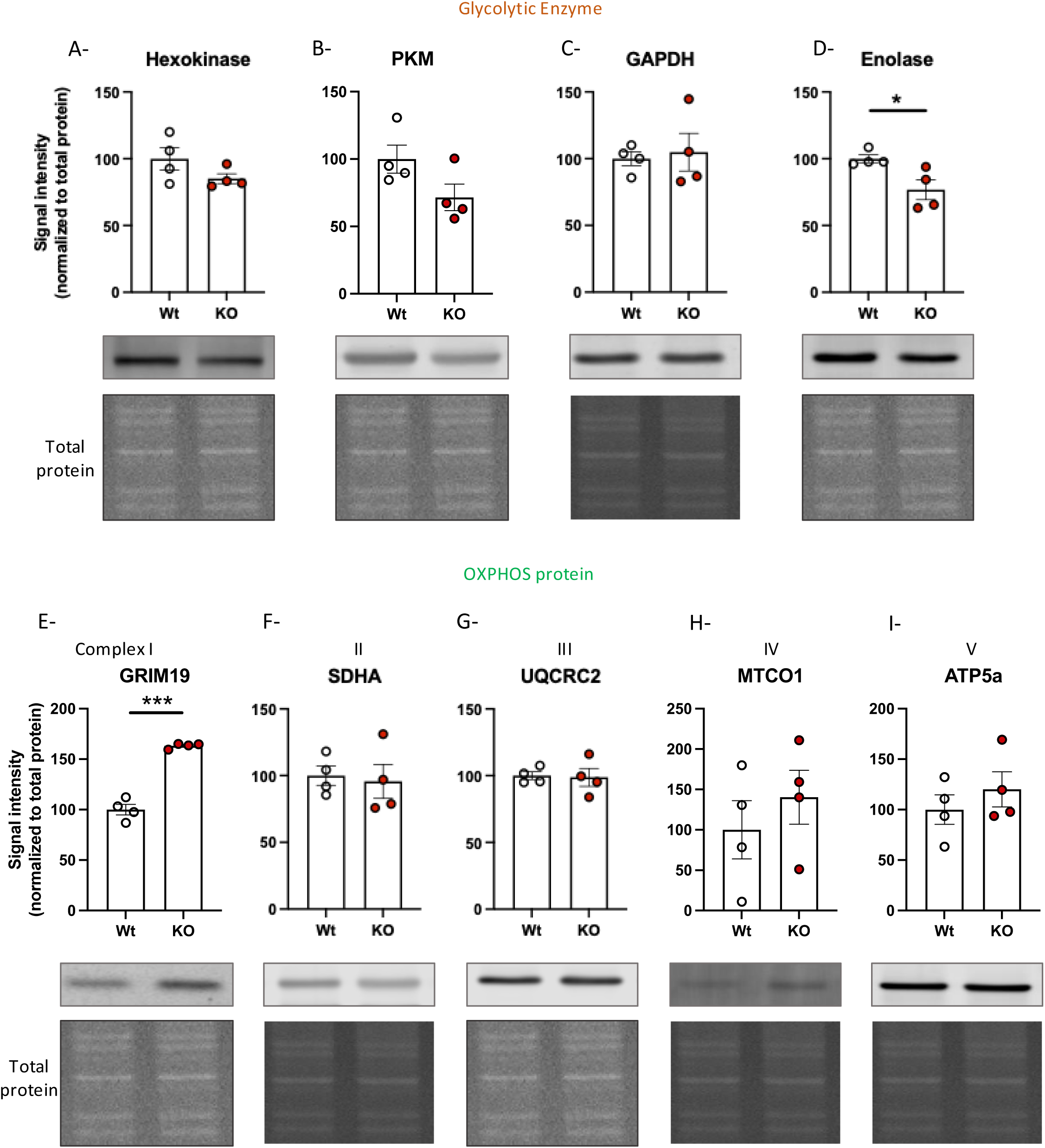
Expression levels of glycolytic enzyme and OXPHOS proteins in N2a cells. A-hexokinase. B-PKM. C-GAPDH. D-Enolase. E-GRIM19. F-SDHA. G-UQCRC2. H-MTCO1. I-ATP5a. J-S1R. A to I (top) Relative signal intensity compared to Wt (bottom) Western blot and total protein staining. Statistical analysis A to I, Student’s t-test (*p < 0.05, ***p < 0.001).

In order to confirm that our findings were caused by the deletion of S1R and not a CRISPR-Cas9 off-target effect, we performed rescue experiments overexpressing S1R in S1R KO N2a cells (Figure S1). First, we overexpressed transiently in S1R KO cells either S1R-GFP or GFP (Figure S1A-D). We observed that the transient expression of S1R slightly rescued the functional glycolytic defect (Figure S1B and C) in a population of 55% GFP positive cells (Figure1S D). To improve a rate of GFP positive cells, we generated a stable cell line overexpressing S1R-GFP and observed increased rescue in KO cells expressing S1R (Figure S1E-H), This suggests that glycolytic activity was correlated with the numbers of S1R-positive cells in the population. Furthermore, we analyzed the protein content in the S1R KO cells with S1R overexpression. We noted that enolase expression seemed to increase back to a normal level (Figure S1I). The concentration of GRIM19 was reduced to a near-physiological level (Figure S1J). We also observed a decrease in the concentration of pyruvate dehydrogenase (PDH) in the S1R KO condition, rescued by the overexpression of S1R-GFP (Figure S1K); however, lactate dehydrogenase (LDH) concentrations were unaffected by the S1R KO condition (Figure S1L). PDH and LDH convert pyruvate, the end-product of glycolysis, into acetyl-CoA in the mitochondria and lactate in the cytosol, respectively.

### Knocking out S1R decreases glycolysis but not oxygen consumption in primary culture of cortical neurons

The N2a cell line was established from mice neuroblastoma, and actively dividing cells have different energy needs different from that of post-mitotic neurons. To confirm the relevance of S1R in primary neurons we repeated the experiments described above on primary cultures of cortical neurons. We first used the sequence of drugs for glycolytic measurement, Rot/AA and then 2-DG. We observed that the primary culture of cortical neurons with S1R KO demonstrated a deficit in basal glycolysis and % PER from glycolysis (Figure 3), consistent with what we observed in N2a cells (Figure 1). Moreover, the S1R KO cortical neurons did not demonstrate a change in oxygen consumption rate (Figure 3A), these results led to an increased mitoOCR/glycoPER (Figure 3B). We also measured the protein concentration of enolase and did not observe any changes (Figure 3C), suggesting that the decrease in enolase concentration observed in the N2a cells models is not essential for the change in glycolytic activity.

**Fig 3.**
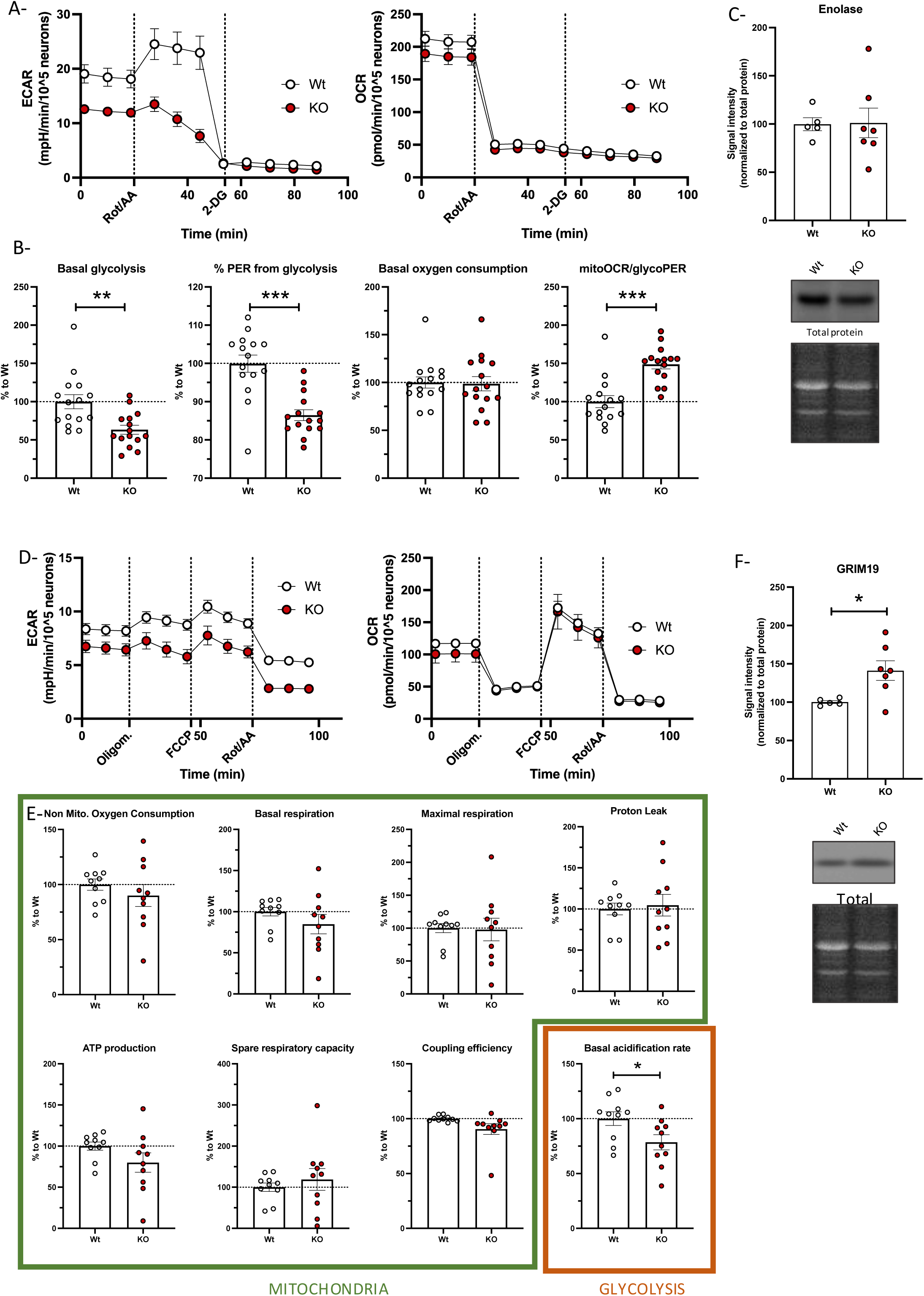
Glycolysis activity and mitochondrial respiration primary culture of cortical neurons. A (left)-Cortical neurons extra-cellular acidification rate (ECAR) measurement with the application of Rotenone and Antimycin A (Rot/AA) and 2-Deoxyglucose (2-DG); A (right)-Simultaneous measurement of cortical neurons oxygen consumption rate (OCR); Neurons from Wt mice in white. Neurons from S1R KO mice in red. B (from left to right)- Basal glycolysis; % proton efflux rate (PER) from glycolysis; Basal oxygen consumption; MitoOCR/glycoPER. C-(top) Enolase signal intensity compared to Wt; (bottom) Western blot and total protein staining. D (left)-Cortical neuronal cell extra-cellular acidification rate (ECAR) measurement with the application of Oligomycin (Oligom.), Carbonyl cyanide-p-trifluoromethoxyphenylhydrazone (FCCP) and Rotenone and Antimycin A (Rot/AA); D (right)-Simultaneous measurement of cortical neurons oxygen consumption rate (OCR); Neurons from Wt mice in white. Neurons from S1R KO mice in red. E (from left to right)-Non mitochondrial oxygen consumption; Basal respiration; Maximal respiration; Proton leak; ATP production; Spare respiratory capacity; Coupling efficiency; Basal acidification rate. F-(top) GRIM19 signal intensity compared to Wt; (Bottom) Western blot and total protein staining. Statistical analysis: Student’s t-test (*p < 0.05, **p < 0.01, ***p < 0.001).

We next measured the same mitochondrial parameters used for N2a cells and found no difference in mitochondria oxygen consumption between S1R KO and Wt cells (Figure 3E). The only change observed in the absence of S1R was the basal acidification rate which was reflective of decreased glycolytic activity (Figure 3D and E). Next, we measured the concentration of GRIM19 in the primary culture of cortical neurons and found increased concentration of GRIM19 in KO condition compared to that of Wt (Figure 3F). These findings were consistent with the results from N2a cells.

Altogether these findings suggest that the S1R deletion impacts glycolysis in both primary cortical neurons and neuronal cell line. Interestingly, in both models, the mitochondria oxygen consumption remained unchanged, although a complex I protein: GRIM19, expression level was increased.

### Knocking out S1R increases NAD^+^/NADH ratio in N2a cells and primary culture of cortical neurons

GRIM19 plays an important role in the complex I which is involved in the regulation of NAD concentrations. Interestingly both glycolysis and mitochondria are impacted and impacting the homeostasis of NAD through the Malate-Aspartate shuttle that allows the regulation of NAD between the matrix of the mitochondria matrix and the cytosol. Therefore, we next chose to investigate NAD metabolism in S1R KO condition.

NAD is a coenzyme present in all living systems. It can be found either in its oxidized or reduced form, NAD^+^ and NADH respectively. We measured the ratio of NAD^+^/NADH in both primary neurons and N2a cells using NAD/NADH-Glo (Figure S2A, Figure 4A-C). We observed in both cortical neurons and in N2a cells that the deletion of S1R induced an unbalanced NAD ratio. Moreover, we found that this ratio was restored to Wt condition levels upon overexpression of S1RGFP in the S1R KO conditions. We also measured the protein expression of SLC25A13, a protein involved in the malate-aspartate shuttle that participates in the regulation of NAD between the mitochondrial matrix and the cytosol, and the concentration of NMNAT-1, a protein responsible for the synthesis of NAD in the nucleus. We did not observe any significant changes in the level of SLC25A13 in KO condition (Figure S2B). However, NMNAT-1 levels were significantly higher in the S1R overexpression condition compared to that of S1R KO and Wt conditions (Figure S2C).

**Fig 4.**
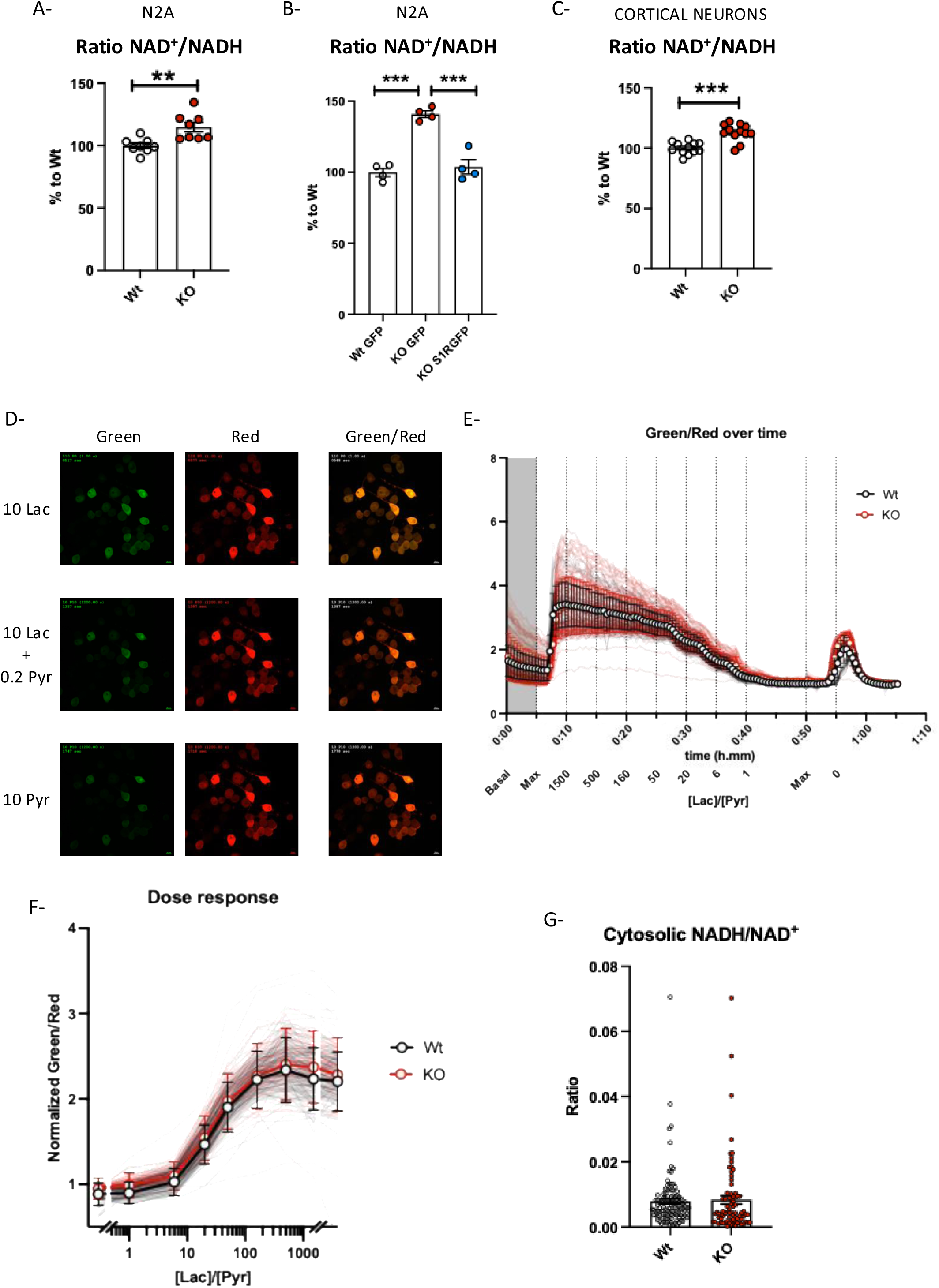
NAD^+^/NADH in N2a cells and primary culture of cortical neurons. A-Wt and S1R KO N2a cell ratio NAD^+^/NADH measured with NAD/NADH-Glo. B-Wt and S1R KO N2a cells transfected with either GFP or S1RGFP plasmid, ratio of NAD^+^/NADH measured with NAD/NADH-Glo; C-Wt and S1R KO primary culture of cortical neuron ratio of NAD^+^/NADH measured with NAD/NADH-Glo. D-Pictures representing Wt N2a cells expressing Peredox with different concentrations of extracellular Lactate and Pyruvate. Green channel represents the Peredox sensor binding NADH and red channel represents the total amount of Peredox expressed in the cell. The ratio of green/ged represents the amount of Peredox binding NADH over the total quantity of Peredox in the cell. E-Representation of the Green/Red ratio over time with the application of a different ratio of [Lac]/[Pyr]. Wt N2a in white. S1R KO N2a in red. F- Dose response of green/red ratio over the [Lac]/[Pyr]. G- Cytosolic NADH/NAD^+^ of Wt and S1R KO N2a cells. Statistical analysis: Student’s t-test (**p < 0.01, ***p < 0.001).

As mentioned earlier in the manuscript the concentrations of NAD^+^ and NADH are subject to active transport among the different organelles in the cell. Particularly in the nucleus, mitochondria and cytosol, the concentrations of NAD forms could vary. The NAD/NADH-Glo assays the concentrations in whole cells lysates. In order to understand how S1R alters the NAD metabolism, we used Peredox, [49, 54] a NAD^+^/NADH sensor specific to the cytosol (Figure 4D-G). Following the protocol described by Hung *et al.* [49, 50] we observed the ratio of NAD^+^/NADH with various concentrations of lactate and pyruvate between Wt and S1R KO N2a cells (Figure 4D, E and Figure S2D). We established a dose response of known concentration of extracellular lactate and pyruvate over the fluorescent intensity of the sensor, then reported the fluorescent intensity in basal condition (without application of lactate or pyruvate) to the dose response. We then calculated the equivalent of NAD^+^/NADH ratio for the corresponding fluorescent intensity (Figure 4G). We did not observe any difference in the NAD^+^/NADH ratio between Wt and S1R KO conditions. This suggests that the whole cell changes in NAD/NADH ratio observed with NAD/NADH-Glo must involve intra-organelles concentrations of NAD.

### Changes in glucose uptake in S1R KO condition

Like NAD^+^ and NADH, glucose is a key substrate for glycolysis. Our data suggest there is a decreased glycolysis with no correlated decrease in glycolytic enzymes. Thus, we next investigated on the ability of different neuron models to import glucose from extracellular media into the cytosol. Using Glucose Uptake-Glo, we observed increased glucose uptake in S1R KO N2a cells; this increased glucose uptake was rescued by the overexpression of S1R (Figure 5A and B). However, when using the same assay on a primary culture of cortical neurons, we observed no differences between Wt and S1R KO cells (Figure 5C). The Glucose Uptake-Glo assay relies on the intake of exogenous 2-deoxyglucose and its metabolization by hexokinase. We did not observe any significant change in hexokinase concentration in S1R KO condition (Figure 2A), suggesting the nature of the N2a could explain the difference in results. We investigated further by using [^18F^]FDG PET-scan in Wt and S1R KO mice (Figure 5). We observed at different time points the regional brain distribution of [^18F^]FDG 30min after its injection on Wt and S1R KO mice (Figure 5D). Mice with S1R KO at 6- and 9-months of age had decreased ^18^FDG uptake compared to that of Wt (Figure 5E), suggesting the possibility that S1R plays a role in brain glucose uptake as a part of the mechanisms on how S1R regulates glycolytic machinery.

**Fig 5.**
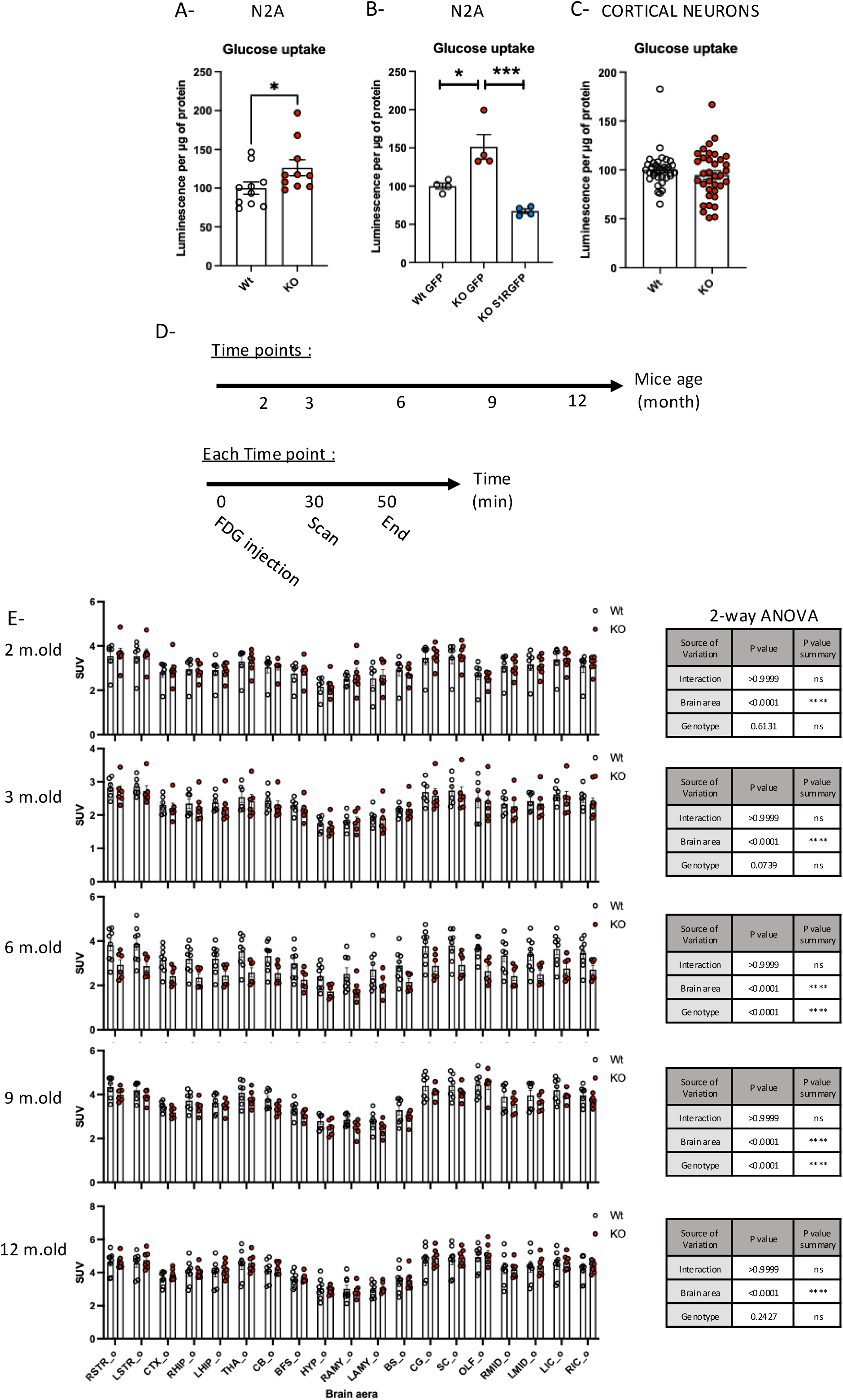
Glucose uptake in S1R KO neuronal models. A-Wt and S1R KO N2a cell glucose uptake measured with Glucose Uptake-Glo. B-Wt and S1R KO N2a cells transfected with either GFP or S1RGFP plasmid; glucose uptake measured with Glucose Uptake-Glo, C-Wt and S1R KO primary culture of cortical neuron glucose uptake measured with Glucose Uptake-Glo, D-PET-Scan ^18^FDG experimental design. E (right)- PET-Scan ^18^FDG results, Standardized uptake value (SUV) per brain area; E (left) 2-way ANOVA for each time point. Statistical analysis for A-C: Student’s t-test (*p < 0.05, ***p < 0.001), for E: 2-way ANOVA (****p < 0.0001).

### Restoring complex I protein level rescues S1R KO phenotype

Our results show that S1R impacts different components of energy metabolism; however, the process by which S1R deletion induced these phenotypes is unclear. S1R is known to stabilize IP3R3 and promote the efflux of calcium from the ER into the mitochondria [1], known to impact the TCA cycle and OxPhos that metabolize NAD [27, 28, 55]. Therefore, we hypothesized that the lack of S1R could alters the mitochondria NAD^+^/NADH ratio and thus impact the glycolysis through alteration of IP3R3. To test this hypothesis, we overexpressed IP3R3 in S1R KO N2a cells (Figure 6A-J). We first ensured that the cell population was expressing turbo GFP tagged IP3R3 (IP3tGFP) or the turbo GFP alone (tGFP). We then tested Wt tGFP, S1R KO tGFP and KO IP3tGFP using Seahorse assays (Figure 6A-C). IP3tGFP overexpression was unable to rescue the glycolytic deficit induced in S1R KO N2a cells when compared to Wt tGFP cells (Figure 6D and E). Interestingly the overexpression of IP3tGFP does not seem to alter the oxygen consumption (Figure 6F and G) nor was unable to rescue the elevated GRIM19 levels in S1R KO N2a cells (Figure 6H). We conclude that the decrease in IP3R3 is not responsible for the energy metabolism S1R KO phenotypes found in S1R KO cells.

**Fig 6.**
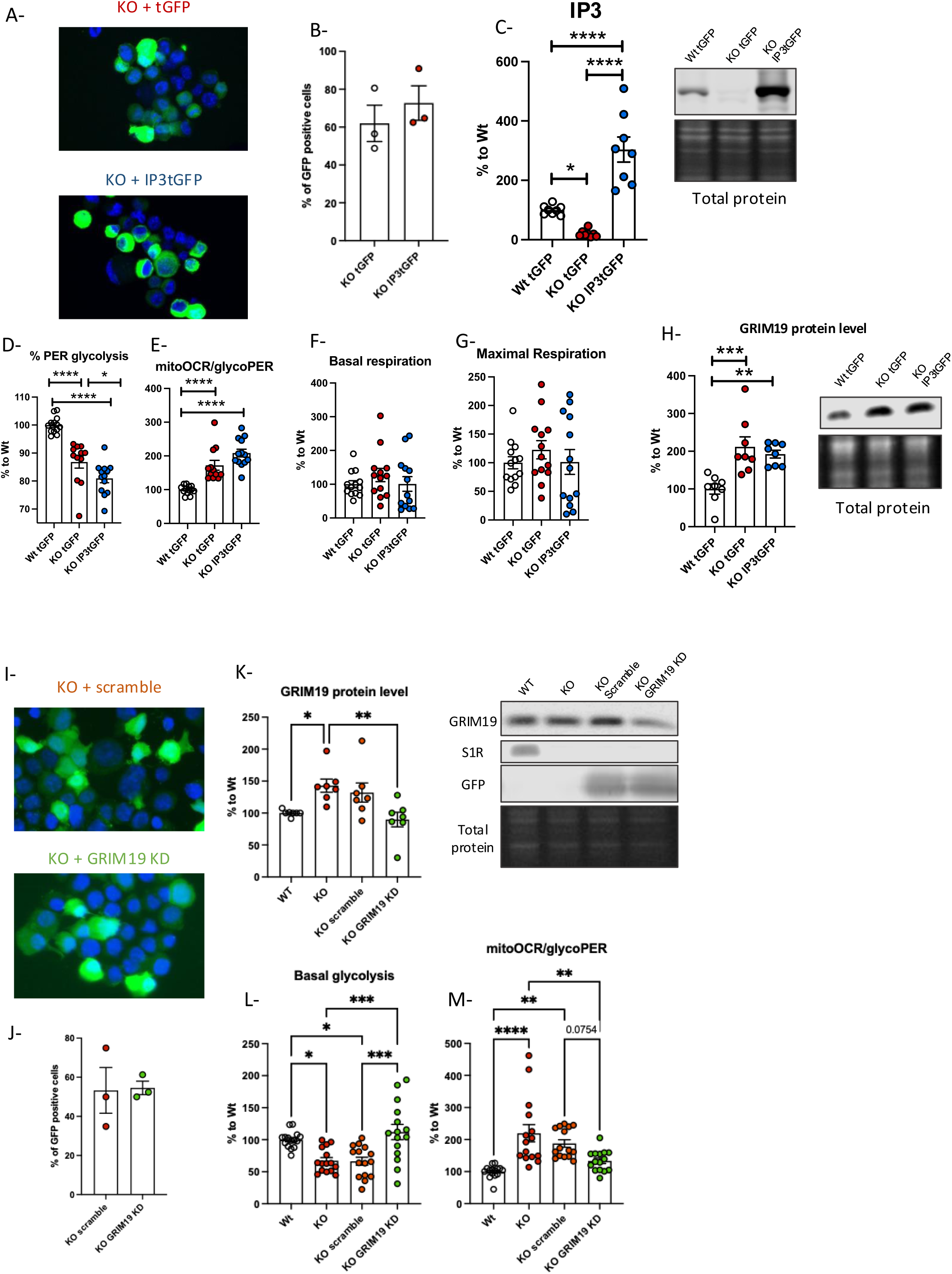
GRIM19 knock-down rescues energy metabolism change induced in S1R KO condition. A- Transfected N2a cells (up : S1R KO tGFP, down : S1R KO IP3tGFP); B- Percentage of GFP-positive cells C-Ip3R3 signal intensity compared to Wt tGFP; D-%PER glycolysis; E-MitoOCR/glycoPER; F- Basal respiration; G-Maximal respiration; H- (left) GRIM19 signal intensity compared to Wt tGFP; (right) Western blot and total protein staining; I-Transfected N2a cells (up : S1R KO scramble, down : S1R KO GRIM19 KD); J- Percentage of GFP-positive cells; K-GRIM19 signal intensity compared to Wt ; L-Basal glycolysis; M-MitoOCR/glycoPER. Statistical analysis, One-way ANOVA followed by multiple comparisons (*p < 0.05, **p < 0.01, ***p < 0.001, ****p < 0.0001)

Based on our finding on consistent increase of GRIM19 in the various S1R KO models, we next investigated whether GRIM19 up-regulation plays a role in the energy metabolism changes induced by the deletion of S1R, or whether GRIM19 is a side target of the energy metabolism changes (Figure 6I-M). We knocked-down GRIM19 in S1R KO cells to return levels to Wt cell levels (Figure 6K-M). We used Seahorse to compare the glycolytic activity of Wt cells, S1R KO cells and S1R KO cells transfected with a scramble, and S1R KO cells with a plasmid knocking down GRIM19 (Figure 6K). The knockdown of GRIM19 reversed the glycolytic deficit phenotype induced in S1R KO cells (Figure 6L and 6M).

Previous publications have suggested that S1R may alters the transcription, translation, and stabilization of different genes, mRNAs and proteins, respectively, by affecting transcription factors [56, 57] and binding to mRNAs [58]. We therefore tested whether the deletion of S1R altered GRIM19 at the transcriptional level by measuring *Grim19* mRNA with qPCR. We did not observe any significant changes in the *Grim19* expression between Wt and S1R KO both in N2a cells and primary cortical neurons (Figure S4A and S4B). We also checked the translational level of *Grim19* mRNA using polysome profiling but found no difference between Wt and S1R KO N2a cells (Figure S4C). Altogether these results suggest that observed S1R KO decreased-glycolysis phenotype is dependent on increased GRIM19 expression, highlighting the connection between S1R and complex I of the mitochondria. Future experiments may be able to reveal the nature of the interaction between S1R and GRIM19.

## Discussion

In this manuscript we described for the first time in neuronal models the relevance of S1R to glycolysis and GRIM19 regulation (Figure 7). We first observed in N2a cells, that have a genetic deletion of S1R, a significantly decreased glycolytic activity without altered mitochondrial functions. These results were mirrored in mouse primary culture of cortical neurons, where we also identified a decreased glycolytic activity in S1R KO neurons without change in mitochondria function. In both N2a cells and cortical neurons we found increased GRIM19, a protein belonging to the complex I of the OxPhos that is involved in NAD metabolism. We then measured the NAD^+^/NADH ratio in the whole cell and cytosolic compartments of N2a cells. Despite seeing an increase in the ratio of NAD^+^/NADH in S1R KO whole-cell lysate, we did not see any changes in the cytosolic compartment. This suggests that the deletion of S1R may impact the concentration of NAD^+^ and NADH stock in other subcellular compartments. We then measured glucose uptake of Wt and S1R KO mouse brains *in-vivo* using FDG-PET. S1R KO mice presented age-dependent decreases in glucose uptake in brain. Finally, to better understand the mechanism by which S1R affects glycolysis, we sought to rescue the S1R KO cells decreased glycolytic activity by separately overexpressing IP3R3 and knocking down GRIM19, which is upregulated in S1R KO cells. Overexpression of IP3R3 did not rescue the glycolytic deficit phenotype observed in S1R KO cells whereas knocking-down GRIM19 significantly did rescue the phenotype.

**Fig 7.**
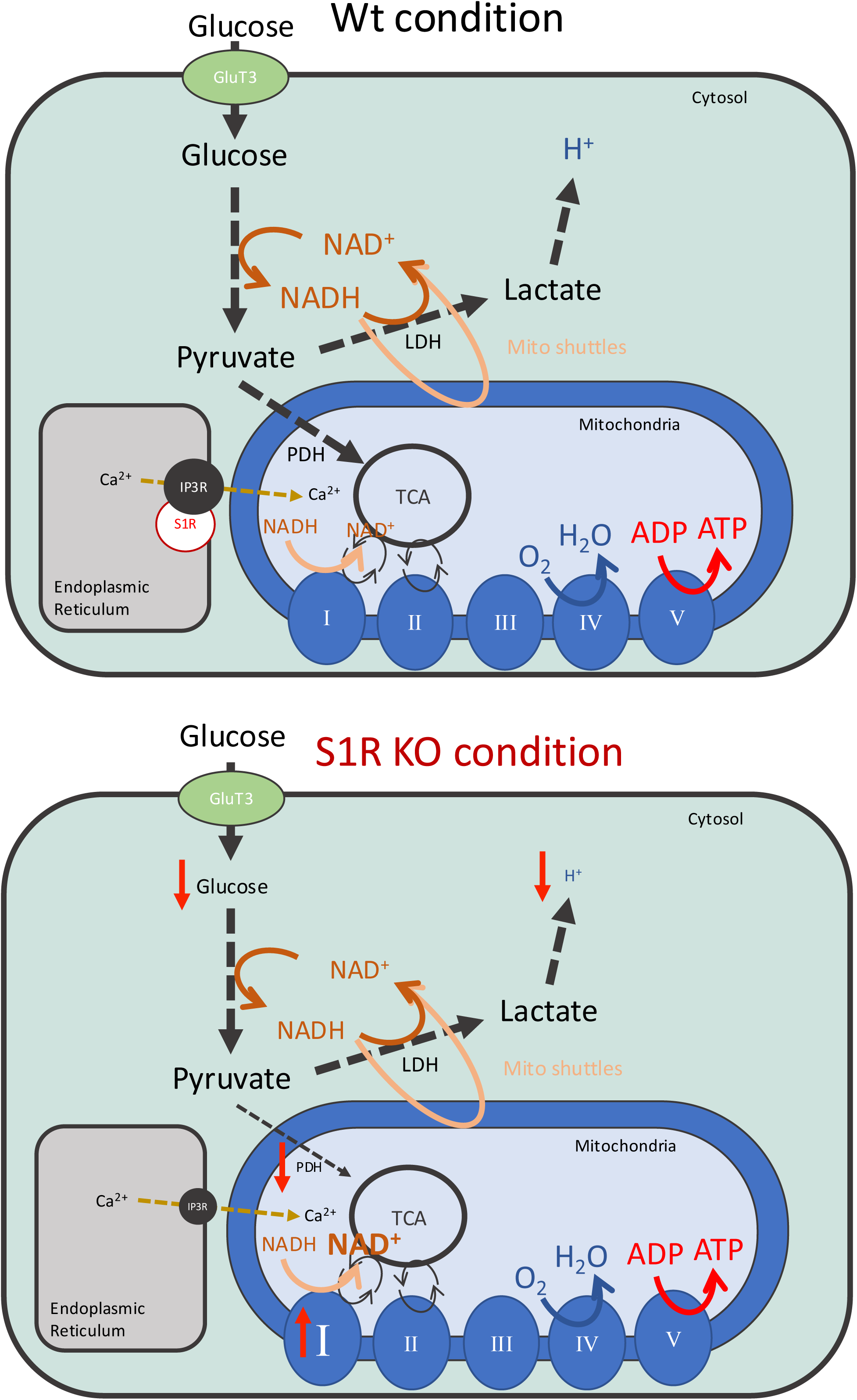
Schem of energy metabolism in Wt and S1R KO condition. Schema highlighting the connection between mitochondrial processes and glycolysis through the NADH/NAD ratio. The role of Complex I of the oxidative phosphorylation chain is included. Summary of the consequences of S1R deletion on the overall energy metabolism and Complex I.

Often under considered over mitochondria that produce more ATP, glycolysis is fundamental for energy metabolism and is particularly important and singular in brain tissues. Neurons produce pyruvate via glycolysis following glucose uptake; however, another mechanism for pyruvate production in neurons has been hypothesized. Several decades ago, it was suggested that neuronal pyruvate can also be produced from lactate transported from supportive astrocytes. This is called the astrocyte to neuron lactate shuttle (ANLS) hypothesis [40, 59]. It is uncertain whether glycolysis or the ANLS is the predominant mechanism for neurons to produce pyruvate [60]. Recently with the help of modern tools and technologies, studies have shown that neurons rely on their own glycolysis to produce pyruvate. Moreover, during neuronal activation, the stoichiometry between glucose and oxygen consumption changes in favor of glycolysis, a less efficient but quicker production of ATP [61–64]. Thus, leads us to think that the relevance of S1R on glycolysis could be particularly critical in neuronal context.

Across the different S1R KO neuronal models used in this study, we consistently observed a decrease in glycolytic activity and an increase in GRIM19 protein level. However, the impact of S1R KO on glucose uptake, and glycolytic enzyme concentrations varies among the models. We propose that this could be explained by the fact that energy metabolism is subject to different constraint between N2a cells, primary neurons and intact brain [65, 66]. Furthermore, knocking down of GRIM19 is sufficient to restore the decrease in glycolysis in S1R KO cells. Therefore, the GRIM19 dysregulation-caused glycolytic reduction may be a direct consequence of S1R deletion whereas changes in glucose uptake and glycolytic enzyme expressions may reflect model-specific compensatory mechanisms. How S1R affects GRIM19 expression remains unknown, although our data show that steady state levels and translation initiation of GRIM19 mRNA are not affected by S1R removal.

S1R is known to promote calcium entry into the mitochondria through stabilizing IP3R3 in the ER membrane. However, the extent to which S1R-facilitated stabilization of IP3R3 affects mitochondrial activity is less known. In this study we did not observe any effect on mitochondrial oxygen consumption following the constitutive deletion of S1R. Bernard-Marissal *et al.* observed that in S1R KO condition the morphology and transport of mitochondria are altered; however they did not assess mitochondria oxygen consumption or ATP production [67]. Abdullah *et al.* observed in the cardiomyocytes of S1R KO mice a decrease in basal mitochondrial oxygen consumption compared to that of Wt. Their results found an increase of complex I protein but a decrease in its activity [68]. In contrast, Motawe. *et al.* did not observe a significant change in mitochondrial oxygen consumption in S1R-knockdown compared to that of a control condition [47]. Another study using two different cell lines found that S1R antagonist does not affect mitochondria bioenergetic processes [22]. It has to be noted that S1R endogenous activation is not well understood and may vary depending on the cellular model, observation made in a cellular context where S1R is not endogenously activated could explain the lack of phenotypes in S1R KO conditions, KD conditions or observed with S1R pharmacological inactivation.

GRIM19 is part of the complex I (also known as NADH ubiquinone oxidoreductase) of the OxPhos chain [55]. From a molecular point of view GRIM19 (also known as NDUFA13) is a nuclear-encoded protein that contributes to the structural integrity of the complex I’s proton channel; therefore, GRIM19 is directly involved in the OxPhos proton transfer [69]. Complex I, along with the complexes III and IV conducts proton transfer from the mitochondrial matrix into the intermembrane space [53]. Interestingly, complex I does not participate directly in mitochondrial oxygen consumption (the role of complex IV) nor in the production of ATP (the role of complex V). Our results show that S1R deletion does not seems to impact the mitochondrial oxygen consumption of mitochondria, across different neuronal models; this is consistent with our finding that concentrations of complexes V and IV remained constant between S1R KO and Wt cells. Interestingly in mouse mitochondrion extracts Goguadze *et al.* observed that pharmacological activation of S1R under physiological conditions increases complex I activity without affecting respiration rate [52]. This implies that genetical or pharmacological manipulation of S1R can alter complex I enough to impact NAD metabolism, without having a significant overall effect on the OxPhos chain.

NAD metabolism is the key link between mitochondrial activity and glycolysis. Communication between these two cellular mechanisms is especially important when the cell faces cellular stress [70–72] and with aging neurons where energy metabolism undergoes drastic changes [37, 73–75]. S1R is a proven therapeutic target in various neurodegenerative disorders such as Alzheimer’s Disease, Huntington’s Disease and Amyotrophic Lateral Sclerosis. Our findings suggest the possibility of extending the therapeutic potential of S1R to pathological conditions with disordered energy metabolism such as cancer and aging.

## Glossary

[^18F^]FDG: [^18^F]fluorodeoxyglucose
2-DG: 2-deoxyglucose
AA: antimycin-A
ECAR: extracellular acidification rate
ER: endoplasmic reticulum
FCCP: carbonyl cyanide p-(trifluoromethoxy)phenylhydrazone
GAPDH: glyceraldehyde-3-phosphate dehydrogenase
GFP: green fluorescence protein
glycoPER: PER from glycolysis
GRIM19: NADH dehydrogenase [ubiquinone] 1 alpha subcomplex subunit 13
GRP75: glucose-regulated protein 75
IP3: inositol 1,4,5-trisphosphate receptor
IP3R3: IP3 receptor type3
KO: knocking-out
LDH: lactate dehydrogenase
MAM: mitochondria-associated ER membrane
MCU: mitochondrial calcium uniporter
mitoOCR: OCR from mitochondria
N2a: Neuro2a
NAD: nicotinamide adenine dinucleotide
NAD^+^: oxidized NAD
NADH: reduced NAD
OCR: oxygen consumption rate
OxPhos: oxidative phosphorylation
PDH: pyruvate dehydrogenase
PER: proton efflux rate
PET: positron emission tomography
qPCR: quantitative polymerase chain reaction
Rot: rotenone
S1R: sigma-1 receptor
shRNA: short hairpin RNA
siRNA: small interfering RNA
tGFP: turbo GFP
TOM70: translocase of outer mitochondrial membrane protein 70
VDAC1: voltage-dependent anion channel 1
Wt: Wild-type

## Author contributions

Conceptualization: SC; methodology: SC, YY, YK, HW, GP; formal analysis and investigation: SC, YY, IG, JH, JG ; writing and original draft preparation: SC; review and editing: SC, YY, IG, YK, JH, JG, HW, AL, MM, BH, and T-PS; funding acquisition: AL, MM, BH, T-PS. All authors read and approved the final manuscript.

## Acknowledgement

We would like to thank Genie Han for language editing. We also would like to thank the laboratory of Genome Integrity: Flow Cytometry Core at the National Cancer institute. This project is supported by the Intramural Research Programs of the National Institute on Drug Abuse (*e.g.* ZIADA000069), National Cancer Institute, and National Institute on Aging.

---

**Supplementary Fig 1.**
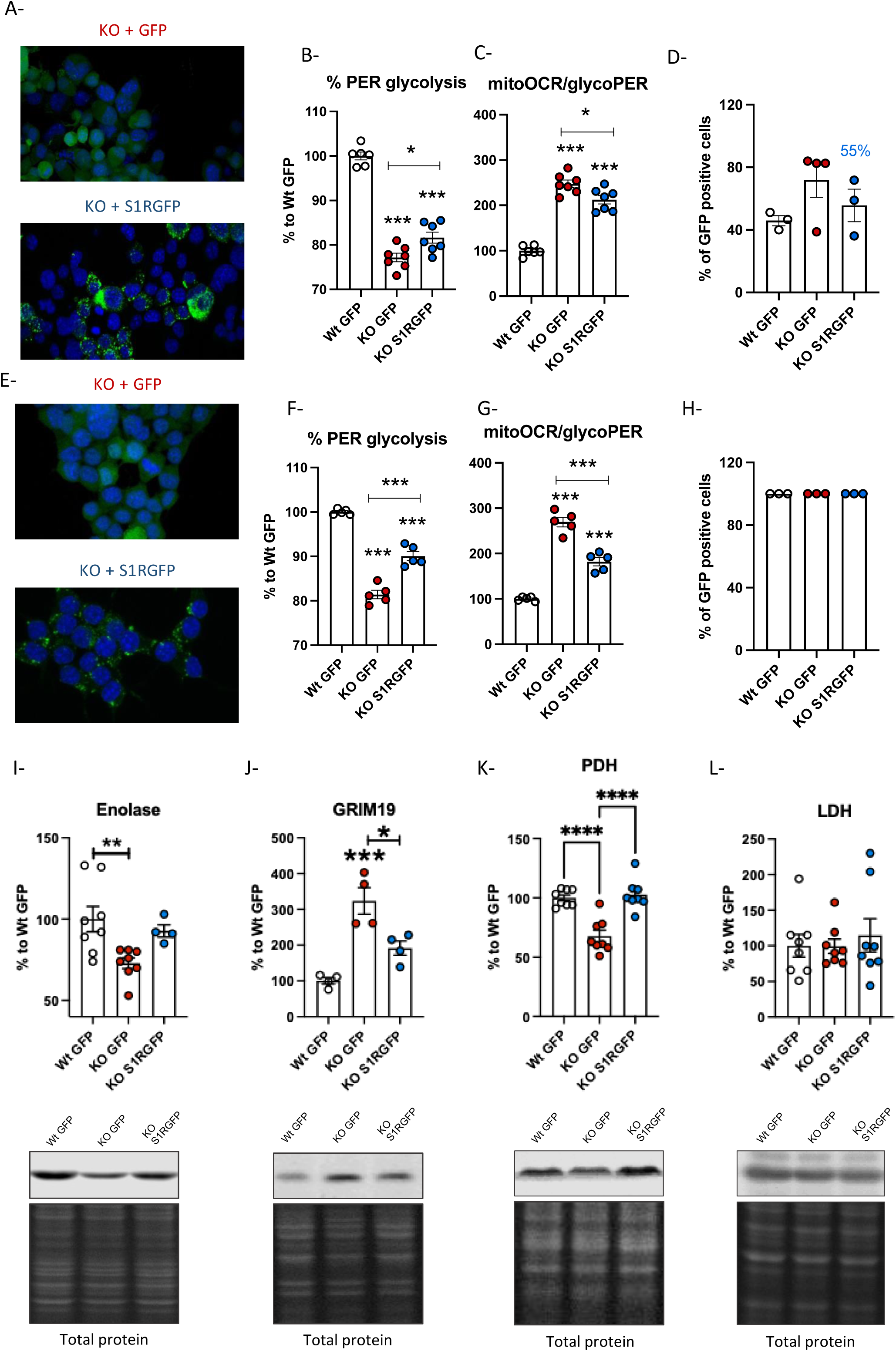
S1R overexpression rescues energy metabolism change. A- Transiently transfected N2a cells (up : KO GFP, down: KO S1RGFP). B-%PER glycolysis. C-MitoOCR/glycoPER. D- Percentage of positive cells. E-Stabilized S1R-overexpressing N2a cells (up : KO GFP, down : KO S1RGFP);. F-%PER glycolysis. G-MitoOCR/glycoPER. H- Percentage of positive cells. I- (top) Enolase signal intensity compared to Wt; (bottom) Western blot and total protein staining. J- (top) GRIM19 signal intensity compared to Wt; (bottom) Western blot and total protein staining. K- (top) PDH signal intensity compared to Wt; (bottom) Western blot and total protein staining. L- (top) LDH signal intensity compared to Wt; (bottom) Western blot and total protein staining. Statistical analysis B, C, F, G, I, J, K and L One-way ANOVA followed by multiple comparisons (*p < 0.05, **p < 0.01, ***p < 0.001)

**Supplementary Fig 2.**
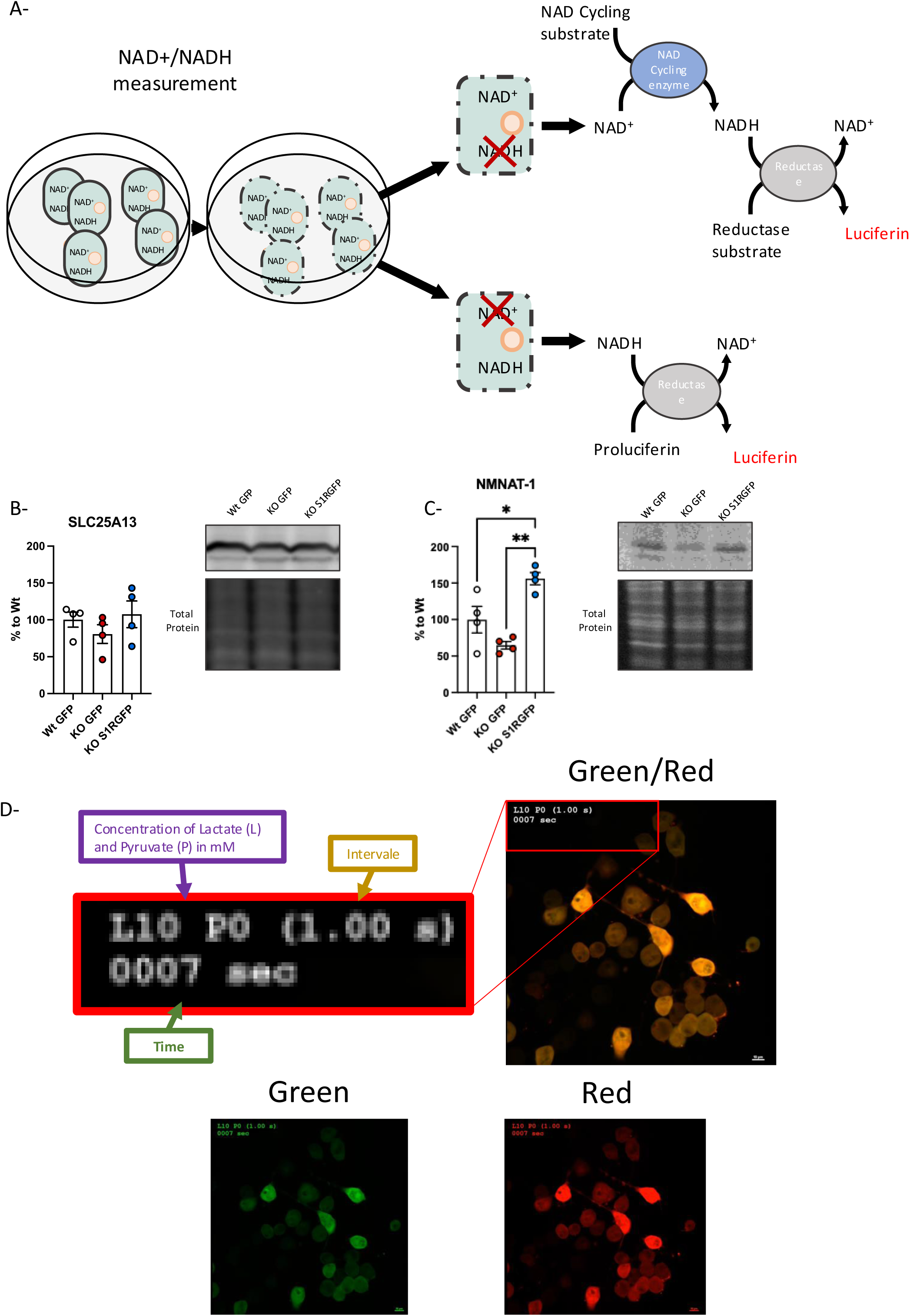
NAD/NADH-Glo™ and Peredox. A-Schema highlighting the NAD^+^/NADH uptake assay and its use with N2a cells and primary cortical neurons. B- Signal intensity of SLC25A13 compared to that of Wt protein level. C- Signal intensity of NMNAT3 compared to that of Wt protein level. D- Video of Peredox fluorescent intensity changes with an exogenous application of lactate and pyruvate. Statistical analysis was one-way ANOVA followed by multiple comparisons (*p < 0.05)

**Supplementary Fig 3.**
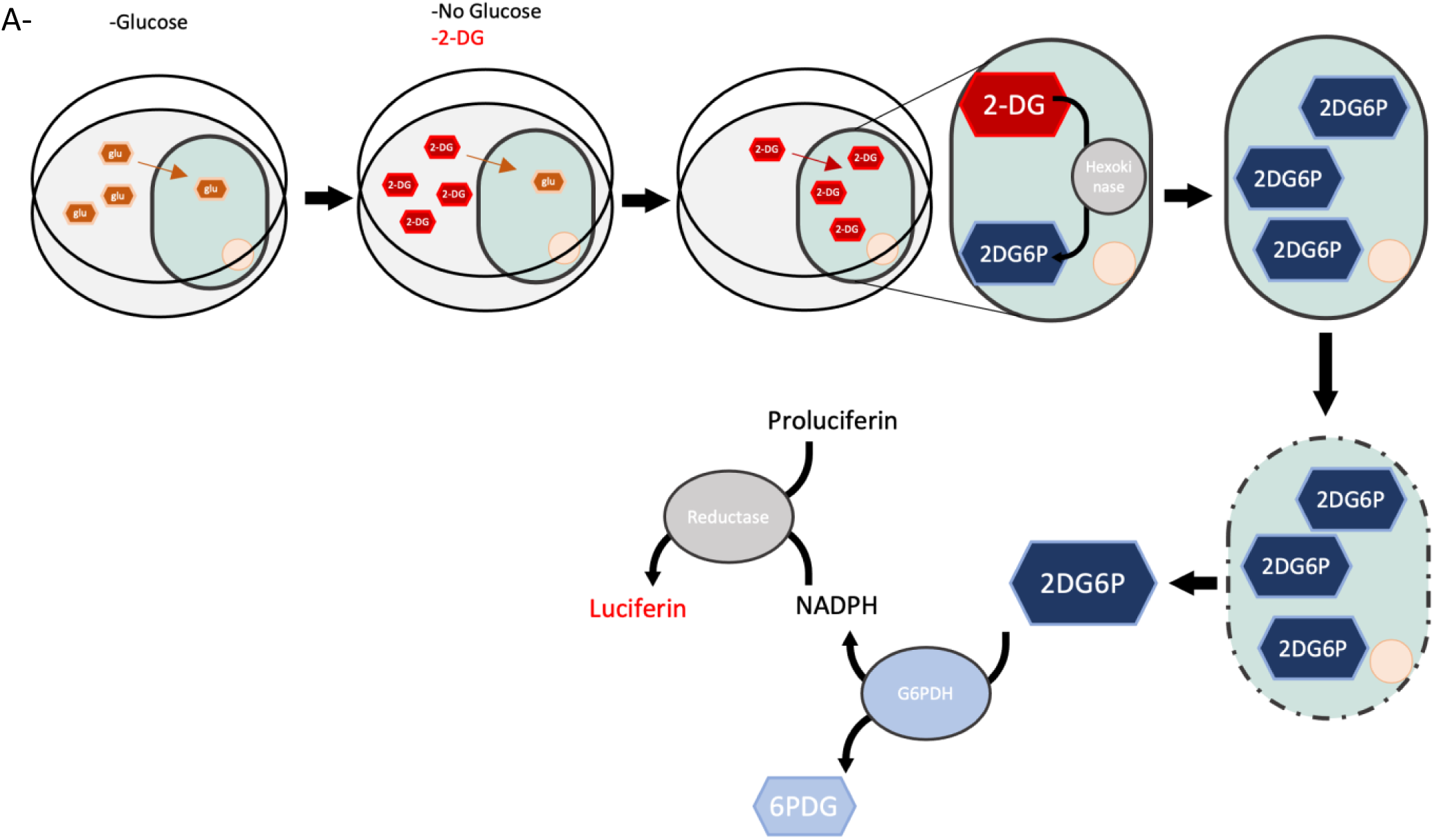
Glucose Uptake-Glo™. A-Schema highlighting the Glucose Uptake assay and its use with N2a cells.

**Supplementary Fig. 4.**
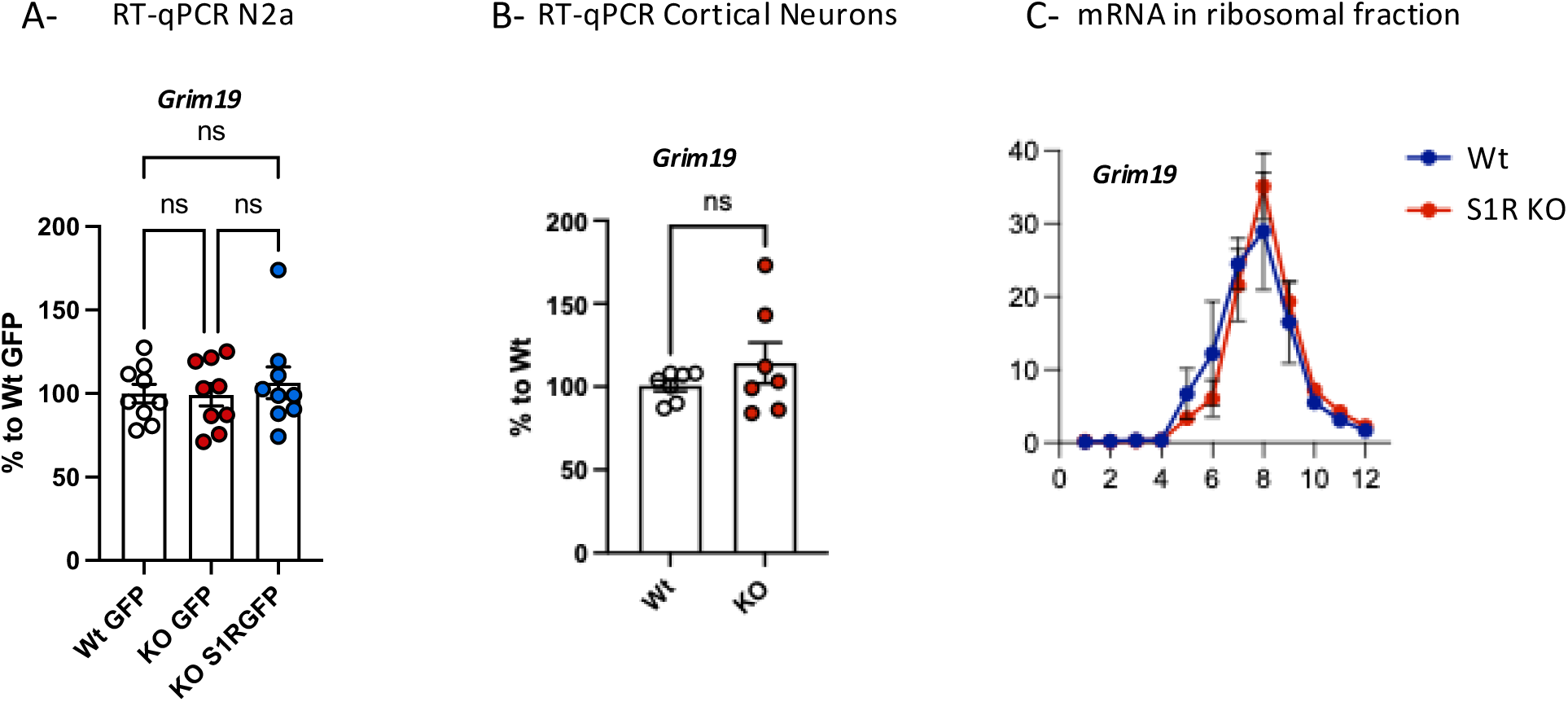
RT-qPCR and mRNA in ribosomal fractions. A- Relative *Grim19* mRNA expression in A- N2a cells and in B- primary culture of cortical neurons. C- *Grim19* mRNA distribution across ribosomal fractions. Statistical analysis for A and B: Student’s t-test.

