## Appendix for "Sigma-1 Receptor Promotes Glycolysis in Neuronal Systems by Suppressing GRIM19"

Appendix 1: Antibody and plasmids list

| Antibody target  (primary antibody) | Company | Ref | Concentration |
| --- | --- | --- | --- |
| Sigma-1 receptor | proteintech | 15168-1-AP | WB 1/500 |
| Hexokinase | Proteintech | 22029-1-AP | WB 1/2000 |
| PKM | cell signaling | C103A3 | WB 1/1000 |
| GAPDH | cell signaling | 5174S | WB 1/1000 |
| Enolase | Santa cruz | sc-271384 | WB 1/500 |
| GRIM19 | proteintech | 10986-1-AP | WB 1/500 |
| SDHA | proteintech | 14865-1-AP | WB 1/800 |
| UQCRC2 | proteintech | 14742-1-AP | WB 1/1000 |
| MTCO1 | Invitrogen | 459600 | WB 1/500 |
| ATP5A1 | proteintech | 14676-1-AP | WB 1/2000 |
| PDH | cell signaling | 3205S | WB 1/1000 |
| LDHA | cell signaling | 2012S | WB 1/1000 |
| Antibody target and fluorophore  (secondary antibody) | **Company** | **Ref** | **Concentration** |
| anti-mouse IgG 800 | LiCor | 926-32210 | WB 1/10000 |
| anti-rabbit IgG 680 | LiCor | 926-68071 | WB 1/10000 |
| Plasmids | **Company** | **Ref** |  |
| Itpr3 (NM_080553) Mouse Tagged ORF Clone | Origene | CAT＃: MG225699 |  |
| pCMV6-AC-GFP Mammalian Expression Vector | Origene | CAT＃: PS100010 |  |

Appendix 2: Peredox media

| Ratio [Lac]/[Pyr] | Sodium Lactate concentration [mM] | Sodium Pyruvate concentration [mM] |
| --- | --- | --- |
| Max | [10] | [0] |
| 1500 | [10] | [0.0067] |
| 500 | [10] | [0.02] |
| 160 | [10] | [0.0625] |
| 50 | [10] | [0.2] |
| 20 | [10] | [0.5] |
| 6 | [10] | [1.6] |
| 1 | [10] | [10] |
| 0 | [0] | [10] |
